# Mathematical modelling evolutionarily stable behavior of zooplankton with state constraints

**DOI:** 10.1101/2020.09.29.319079

**Authors:** O. Kuzenkov, E. Ryabova, A. Garcia, O. Kostromina

## Abstract

The purpose of this work is to create mathematical base and software for solving the problem of finding an evolutionarily stable strategy of zooplankton diel vertical migrations and explaining the observed effects in aquatic ecosystems using this software (in particular, in the northeastern part of the Black Sea). An essential feature of this study is the inclusion in the mathematical model of state constraints on the strategy of behavior, which reflect the vertical limited zone of zooplankton habitat. The presence of state constraints creates the main mathematical difficulties for solving the optimal control problem used in the analysis of the model.

The general methodological basis for defining evolutionarily stable behavior is the Darwinian principle “survival of the fittest”. However, it remains a problem to construct a mathematical expression for the fitness function of hereditary elements. The efforts of the authors were aimed at creating a software package that allows predicting the evolutionarily stable behavior of zooplankton based on the actual universal extreme principle. The created software package includes, as a main component, a computational module for solving the set optimal control problem with state constraints.

## 1 Introduction

Currently, methods of computer simulation are widely used to study various phenomena in different subject areas of science, including areas where their application previously seemed impossible. Information technology has long been used in biology (see [1] for a short review). However, they recently began to be used to identify evolutionarily stable hereditary characteristics of living organisms [2–5].

Hereditary traits are called evolutionarily stable if they persist in the population during the struggle for existence in a stable environment [6]. Presence of carriers of such traits is resistant to the introduction into the population of a certain number of mutants with different characteristics. Obviously, knowledge of evolutionarily stable traits is equivalent to knowledge of the biological evolution results for a given population. This, in turn, is crucial for studying the evolution of ecosystems.

The study and modeling of biological evolution is one of the most difficult scientific problems. It requires an integrated approach to adequately describe the physical and chemical factors of the environment and the behavioral reactions of living organisms.

It is clear that an adequate solution to such a complex problem is inconceivable without the non-trivial use of mathematical modeling methods and modern information technology tools. It is necessary to develop a methodology for modeling evolutionary processes and create software packages to support such modeling.

The purpose of this work is to create mathematical base and software for solving the problem of finding an evolutionarily stable strategy of zooplankton diel vertical migrations and explaining the observed effects in aquatic ecosystems using this software (in particular, in the northeastern part of the Black Sea).

The article continues the series of author’s works devoted to modeling zooplankton behavioral strategies and creating a package of applied programs to support such research [2–6]. An essential feature of this work is the inclusion in the mathematical model of state constraints on the strategy of behavior, which represent the most difficult case for the solution of the optimal control problem used to analyze the model.

## 2 Sources review

The phenomenon of diel vertical migrations of aquatic organisms was discovered more than 200 years ago [7]. For the first time in 1817, the explorer Cuvier recorded accumulations of Daphnia in the morning and in the evening near the surface of the water of Lake Geneva and their movement to depth in bright daylight. The first targeted studies of diel vertical migrations of crustaceans were carried out by Forel at Lake Lehmann in 1874 and Weismann at Lake Constance in 1877. Subsequently, it was found that many species of marine and freshwater zooplankton drift long distances during the day, rising at dusk to the water surface and deep down at dawn [7]. Some results of recent observations are given in [2, 8].

The study of the zooplankton behavior is of great importance, since it plays a key role in aquatic ecosystems, is the basis of food chains, makes a significant contribution to carbon exchange and potentially affects the planet’s climate [9]. Diel vertical migration of zooplankton is one of the most significant synchronous movements of biomass on Earth [10].

The phenomenon of diel vertical migrations has been widely studied both empirically and theoretically. However, there is no consensus on how environmental variables affect this movement. For example, changes in light intensity at dusk and dawn, ultraviolet radiation, the size of crustaceans, moonlight, avoidanceof predators, low temperatures, and increased genetic exchange may be key factors.

Studying the behavior of zooplankton is a challenging task for biologists, since such behavior is very diverse even for related species and for different age groups of the same species [2, 7].

To achieve the calculated quantitative prognostic effect, it is inevitable to use mathematical modeling and create the necessary software. Various mathematical models of zooplankton behavior have been considered by a wide range of researchers [11–16].

In particular, diel migrations were considered as an adaptive effect of an organism to environment, as a result of evolution in the struggle for existence [17]. In a number of works, special attention was paid to the construction of specific software [7].

However, the question of the correctness of the mathematical expression of the fitness function remains open. Similar expressions are often subjective and depend on the personal preference of a researcher. Sometimes different approaches predict conflicting behaviors [18, 19]. There is no single software package that would take into account a wide variety of environmental factors and complex relationships between different groups of the population.

The efforts of the authors were aimed to creating a software package that would allow us to predict the evolutionarily stable behavior of zooplankton based on a well-grounded universal extreme principle [6].

## 3 Materials and methods

The general methodological basis for defining evolutionarily stable behavior is the Darwinian principle “survival of the fittest” [20]. However, it remains a problem to construct a mathematical expression for the fitness function of hereditary elements [21]. This problem has been the subject of many studies, starting with the classics of Haldane, Fisher, and Wright [22].

Now there are many different approaches to solving this problem, in particular, the adaptive dynamics approach is widely used [19, 23, 24]. But, perhaps, the most universal solution was proposed by Gorban to derive the fitness function as the average time value of the reproduction rate [25, 26].

This approach was later developed in the works [6, 27, 28, 32], where algorithms for deriving the fitness function depending on the type of the system of competing communities were proposed. This approach would be applied to construct the fitness function in the zooplankton community [2, 29–31]. In the simplest case, the fitness function consists of the energy gain recieved with food, energy expenditures for vertical movements and losses as a result of predation, unfavorable temperature conditions, and hydrogen sulfide concentration.

The parameters of the fitness function can be identified based on the analysis of time series of the abundance of various species and the corresponding selection processes. It is advisable to apply modern technologies of computer neural networks for solving this problem. The technique of such solving was discussed in detail in [3].

If we know the fitness function, we can find an evolutionarily stable strategy of vertical migrations.

This strategy corresponds to the global maximum of this function. To find such a strategy, it is advisable to use the classical methods of the calculus of variations or optimal control [2, 19, 28–30].

Mathematical research and the solution of the optimal control problem are of key importance. Of course, a full consideration of all empirical data features leads to an optimization problem that does not lend itself to an analytical solution. However, as it often happens in mathematical modeling, a simplified formulation of the problem can provide significant help in understanding the problem, allowing for significant analytical research, and sometimes even an exact solution.

Knowledge of such a solution can significantly reduce the range for possible values of the fitness function identified parameters. The solution of such a reduced problem makes it possible to form a training sample using simplified samples, which, nevertheless, retains the essential qualitative features of the original problem.

At the same time, the correct trade-off must be reached between the degree of simplification and maintaining the adequacy of the model. In the previous works of the authors [4, 29, 32], the model was based on a linear-quadratic optimal control problem without phase constraints. The advantage of this model is the ability to obtain an exact analytical solution. But as practice shows, this simplification in some cases is too inadequate.

These circumstances require the restructuring of the created support taking into account the natural phase constraints of the problem.

The developed software package must include, as the main component, a computational module for solving the posed problem of optimal control. But the created software cannot be limited only to the solution of a mathematical optimization problem due to the inaccuracy and approximation of environmental factors data. In addition, the realizable movement of zooplankton always allows random fluctuations. Therefore, it is more important to determine not the exact solution, but the qualitative characteristics of the behavior. The most important qualitative characteristic is the presence or absence of significant movements of zooplankton. Revealing the presence of such characteristics is carried out by computer recognition tools, in particular, using neural network technologies. In this case, the main difficulty is the creation of training samples. The observational stock is too small to obtain the representative training set. Therefore, one has to resort to computer simulations, in which the exact solutions of the optimization problem for different approximated input data play a central role.

A set of exact solutions for various environmental conditions can be used as a training sample to recognize the qualitative characteristics of evolutionarily stable behavior under conditions of incomplete information about the state of the environment [4]. Moreover it can be used to identify the parameters of the fitness function according to the observed process of selecting strategies [3]. The technique for constructing the software package is discussed in detail in [4, 34]. To solve the optimal control problem, the Maple 17 package is also used. In order to test the created software, we used the empirical data given in [2, 8, 7, 18].

## 4 Results

The created software package contains three interconnected complexes. The aim of the first complex is the identification of fitness function on the base of observed selection results. The second complex provides finding evolutionarily stable behavior of zooplankton by numerical solution of the optimal control problem. It solves the problem of the fitness function maximization for different states of environment. As a result, we can form a set of evolutionary stable trajectories corresponding to different environmental factors. It can be used as a training set of comparison samples. The tried complex recognizes the qualitative characteristics of evolutionarily stable strategy on the base of approximately given characteristics of the environment. It uses the training set formed by the second complex.

The input data of software package is the set of functions describing the current state of the environment. Empirical observations confirm that the evolutionary stable movement of plankton is determined by the following environmental factors: spatial distributions across the depth *x* of food, predator density, temperature, radiation level, the overall predator activity during the day, etc.

To find the evolutionary stable strategy, we use the following model of the zooplankton behavior.

Let’s introduce the notation: *x* -vertical coordinate of the zooplankton position; *t* -time of day, ranging from 0 to 1, with 0 corresponding to noon, ½ -midnight, 1 -next noon; *x*(*t*) -population immersion depth depending on the time of day. It is obvious that the function *x*(*t*) must be continuous periodically extended with a period *T* = 1 (one day). This implies the condition *x*(0) = *x*(1). Zooplankton lives in a vertically limited zone: not above the water level and not below the level with a concentration of hydrogen sulfide that is lethal to animals. We introduce the system so that *x* = 0 -determines the level of the upper position of zooplankton, above which it does not rise, *x* = −*C/σ* the level of subsidence below which zooplankton does not live (*C, σ* – positive constants).

We will denote *E*_0_ = *E*_0_(*x*) -energy gain from food intake; *S*_*x*_(*x*) -number of predators depending on depth; *S*_*t*_(*t*) -predator activity during the day; *S* = *S*_*x*_(*x*) *·S*_*t*_(*t*) -losses as a result of predatory extermination; *G* = *G*(*x*) -additional mortality caused by approaching the boundaries of the habitat (water surface or a level with a lethal concentration of hydrogen sulfide); 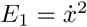 -population losses due to metabolic costs of migration, proportional to the square of the speed of movement. Using these functions, you can compose an energy balance equation that allows you to calculate the energy gain *R* of each individual during the day

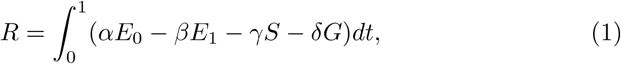

where *α, β, γ, δ* are weight coefficients reflecting the degree of influence of these factors on the fitness of organisms. The calculation of the values of these coefficients is provided by the first software complex based on observational data. It is assumed, that this energy gain is spent on procreation. Then the average time value of the reproduction rate is equal to *R*.

Suppose that the zooplankton population consists of different species, differing from each other in the nature of movement in vertical layers of water. We denote by *v* = *x*(*t*) the hereditary strategy of plankton behavior. The optimal strategy will be the one that maximizes the time average of the reproduction rate corresponding to the strategy.

The problem is to predict the behavior of the zooplankton population after a long time of adaptation on the basis of experimentally recorded input data reflecting the state of the environment.

We can use different approximation of environmental factors.

The simplest approximation is the following linear-quadratic approximation. Let’s take the following approximations: *E*_0_ = *σx* + *C, S*_*x*_(*x*) = *σx* + *C, S*_*t*_(*t*) = cos(2*π*(*t* − *τ*)) + 1 (*τ* is some constant), *G* = (*x* + *C/*(2*σ*))^2^. On the one hand, they represent a kind of approximation to the actually observed data, on the other hand, they make it possible to effectively use analytical methods for solving, which significantly reduces the numerical calculation.

Solving the optimal control problem with state constraints, we obtain that the continuous solution is symmetric with respect to the time instant *t* = 1/2 and on the time interval 0 *≤ t ≤* 1/2 has the form

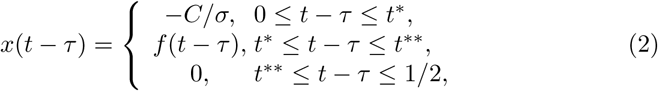

Where

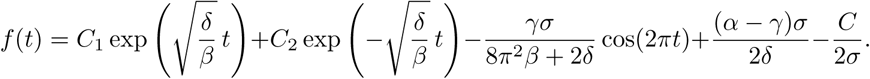

We establish an analytical connection between the constants *C*_1_, *C*_2_ from the condition of the continuity of the solution, numerically choose the pair *C*_1_, *C*_2_ on which the functional (1) reaches its maximum.

For *α* = 0.477, *β* = 2.33203 *·*10^−5^, *δ* = 1.166015 *·*10^−3^, *γ* = 0.521804533, *σ* = 4, *C* = 400 we can represent the environmental factors as shown in Figs. 1–4. Zooplankton trajectory is represented in Fig.5. In this example we got the following values for *C*_1_ and *C*_2_: *C*_1_ = 0.27002 and *C*_2_ = 317.90.

**Fig. 1.**
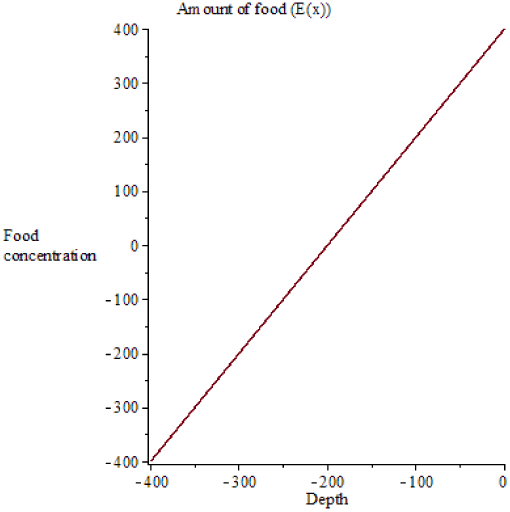
Amount of food *E*_0_(*x*) = *σx*+*C*

**Fig. 2.**
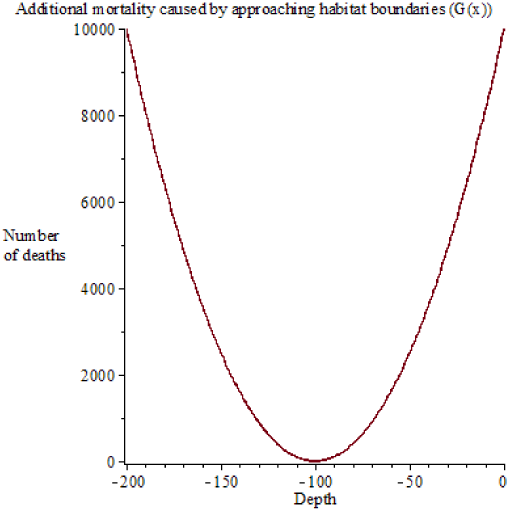
Additional mortality caused by approaching habitat boundaries *G*(*x*) = (*x* + *C/*(2*σ*))^2^

We can test the effectiveness of our model by comparing it with the real observed trajectory of zooplankton migration in Saanich Bay. In Fig. 5, the calculated trajectory of zooplankton movement is compared with the trajectory of the identified vertical movement of zooplankton in Saanich Bay [8]. It can be seen that real and calculated trajectory almost coincide. Therefore even the simplest approximation gives a fairly accurate prediction of evolutionarily stable behavior.

**Fig. 3.**
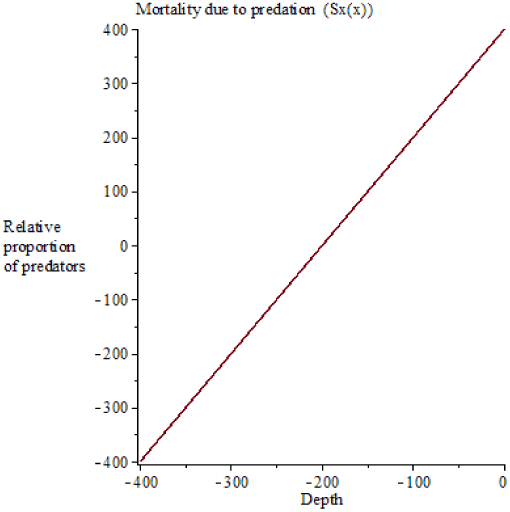
Mortality due to predation *S*_*x*_(*x*) = *σx* + *C*

**Fig. 4.**
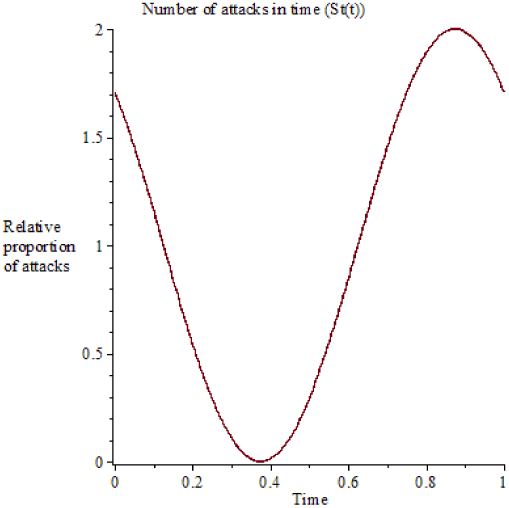
Number of attacks *S*_*t*_(*t*) = *cos*(2*π*(*t* − *τ*)) + 1, *τ* = −3/24

**Fig. 5.**
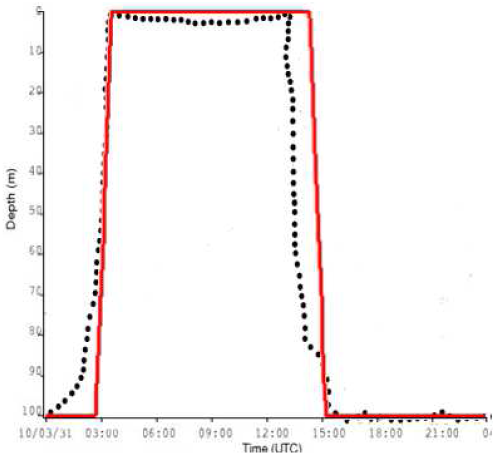
Comparing with the experimental data observed on 01/04/2010 from Saanich Inlet. The red dotted line indicates the path most likely to be followed by zooplankton and the black line is the one obtained by our model for *α* = 0.477, *β* = 2.33203 *·* 10^−5^, *δ* = 1.166015 *·* 10^−3^, *γ* = 0.521804533, *σ* = 4, *C* = 400.

Now, to describe the recorded experimental data of the environment, we will use hyperbolic functions: 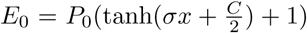, where *P*_0_ represents the maximal phytoplankton density [2], 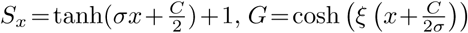. We leave functions *S*_*t*_(*t*), *E*_1_ unchanged. These environmental factors as shown in Figs. 6–9.

**Fig. 6.**
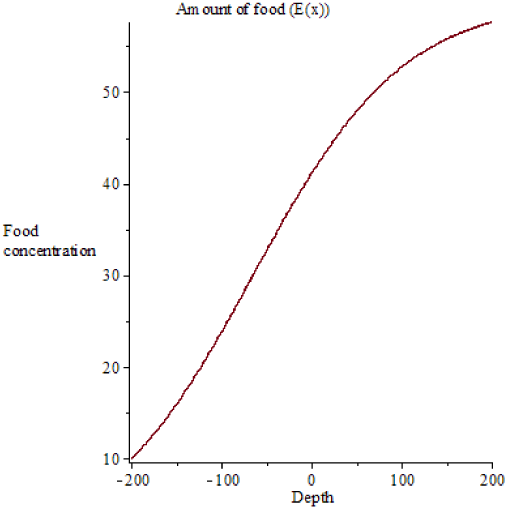
Amount of food 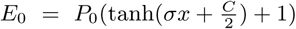

**Fig. 7.**
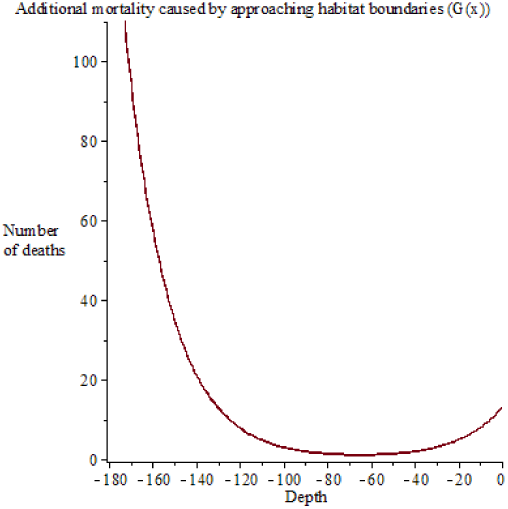
Additional mortality caused by approaching habitat boundaries 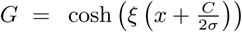

**Fig. 8.**
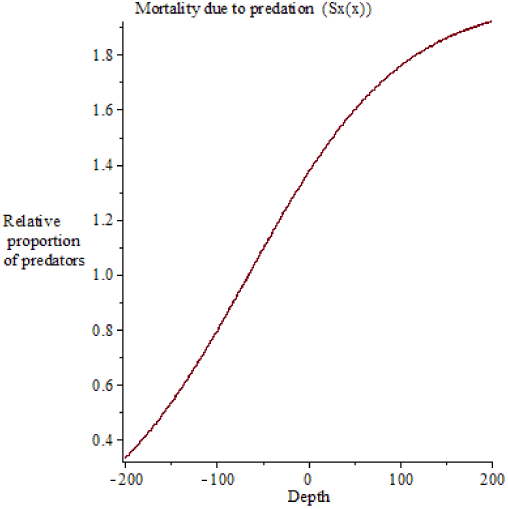
Mortality due to predation 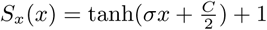

**Fig. 9.**
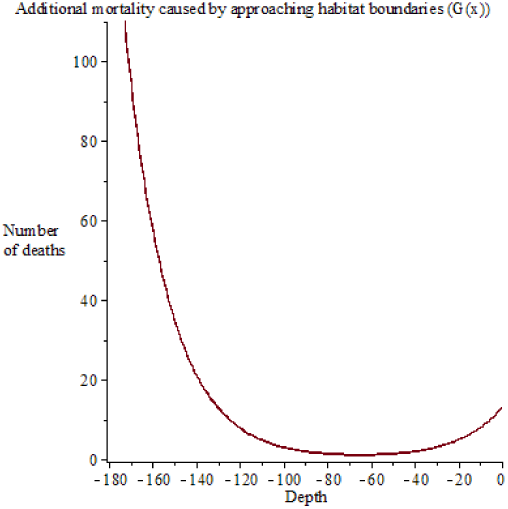
Number of attacks *S*_*t*_(*t*) = *cos*(2*π*(*t* − *τ*)) + 1, *τ* = −0.5

In this case, the trajectory of the zooplankton movement also has the form (2), where the function *f* (*t*) is found numerically.

The graphs of the calculated trajectories of zooplankton for various parameters *α, β, γ, δ, P*_0_, *ξ, σ* and *C* are presented in Figs. 10, 11.

**Fig. 10.**
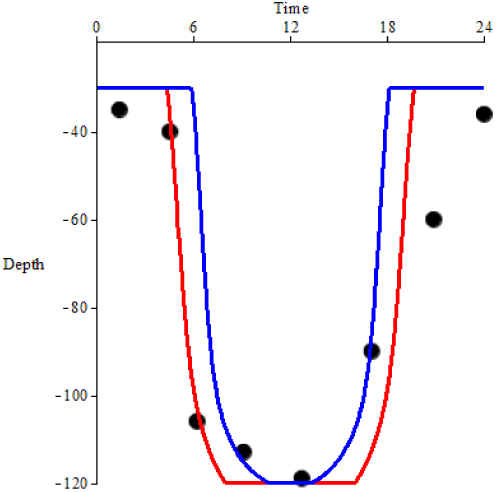
Comparing with the typical dial vertical movement of zooplankton herbivore P. Elongatus observed in Northen Black Sea in summer and collected on 21/06/2009. The black dotted line indicates the path followed by zooplankton, the red line is the one obtained by our model for *α* = 0.04, *β* = 2.8972 *·* 10^−6^, *γ* = 1.6, *δ* = 0.00629944, *P*_0_ = 30, *ξ* = 0.05, *σ* = 0.006 and *C* = 0.72 and the blue line is the one obtained by our model for *α* = 0.08, *β* = 2.8972 *·* 10^−6^, *γ* = 1.6, *δ* = 0.00314972, *P*_0_ = 22.5, *ξ* = 0.06 *σ* = 0.006 and *C* = 0.72

**Fig. 11.**
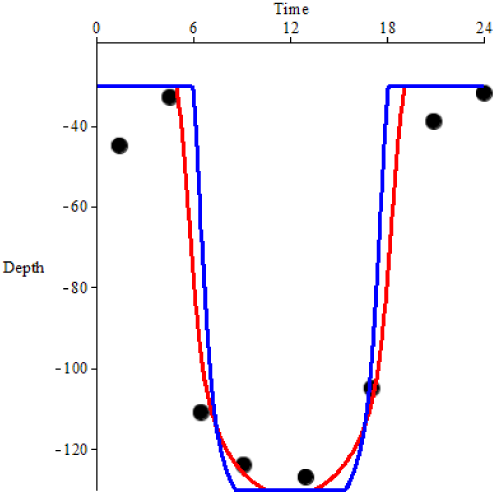
Comparing with the typical dial vertical movement of zooplankton herbivore C. Euxinus observed in Northen Black Sea in summer and collected on 21/06/2009. The black dotted line indicates the path followed by zooplankton, the red line is the one obtained by our model for *α* = 0.05, *β* = 2.8972 *·* 10^−6^, *γ* = 1.6, *δ* = 0.00629944, *P*_0_ = 30, *ξ* = 0.05, *σ* = 0.006 and *C* = 0.78 and the blue line is the one obtained by our model for *α* = 0.08, *β* = 2.8972 *·* 10^−6^, *γ* = 1.6, *δ* = 0.00314972, *P*_0_ = 22.5, *ξ* = 0.05 *σ* = 0.006 and *C* = 0.78

In Figs. 10, 11, the calculated trajectory of zooplankton movement is super-imposed on the graphs of the revealed migratory vertical movement of zooplankton in the northeastern part of the Black Sea [2]. It can be seen that real and calculated trajectory almost coincide. Therefore, the created software allows us to accurately find an evolutionarily stable trajectory.

The obtained results of calculation and computer simulations are included in the set of comparison samples that ensure the work of the third complex.

## 5 Summary

As a result of the work carried out, software was created to solve the problem of finding an evolutionarily stable strategy for diel vertical migrations of zooplankton. To model the behavior of zooplankton, data on the environment are used, such as the presence of food, predators, unfavorable conditions of existence at the boundaries of the habitat, described by linear-quadratic functions or hyperbolic ones. The developed software was tested using observation data in the northeastern part of the Black Sea.

The developed research methodology and software are used in the educational process of the Institute of Information Technologies, Mathematics and Mechanics of Lobachevsky State University of Nizhny Novgorod for the preparation of master’s theses.

The results of this research are used in the educational process of Institute of Information Technology, Mathematics and Mechanics of Lobachevsky State University of Nizhni Novgorod (UNN) within studying of the discipline “Mathematical modeling of selection processes” and for the preparation of Master’s dissertations. It corresponds to the trends of ICT higher education modernization [35, 36] and to the best educational practices of UNN [37–41].

## Notes

### Competing Interest Statement

The authors have declared no competing interest.

